# Uncertainty quantification in subject-specific estimation of local vessel mechanical properties

**DOI:** 10.1101/2021.08.02.454803

**Authors:** Bruno V. Rego, Dar Weiss, Matthew R. Bersi, Jay D. Humphrey

## Abstract

Quantitative estimation of local mechanical properties remains critically important in the ongoing effort to elucidate how blood vessels establish, maintain, or lose mechanical homeostasis. Recent advances based on panoramic digital image correlation (pDIC) have made high-fidelity 3D reconstructions of small-animal (e.g., murine) vessels possible when imaged in a variety of quasi-statically loaded configurations. While we have previously developed and validated inverse modeling approaches to translate pDIC-measured surface deformations into biomechanical metrics of interest, our workflow did not heretofore include a methodology to quantify uncertainties associated with local point estimates of mechanical properties. This limitation has compromised our ability to infer biomechanical properties on a subject-specific basis, such as whether stiffness differs significantly between multiple material locations on the same vessel or whether stiffness differs significantly between multiple vessels at a corresponding material location. In the present study, we have integrated a novel uncertainty quantification and propagation pipeline within our inverse modeling approach, relying on empirical and analytic Bayesian techniques. To demonstrate the approach, we present illustrative results for the ascending thoracic aorta from three mouse models, quantifying uncertainties in constitutive model parameters as well as circumferential and axial tangent stiffness. Our extended workflow not only allows parameter uncertainties to be systematically reported, but also facilitates both subject-specific and group-level statistical analyses of the mechanics of the vessel wall.

## 1 INTRODUCTION

Detailed quantification of the nonlinear, anisotropic mechanical behavior exhibited by blood vessels is essential for understanding the wall mechanics, for informing models of the associated mechanobiology, and for building appropriate fluid-solid-interaction models of the hemodynamics. Such detail is generally possible only for excised specimens subjected to well-defined boundary conditions. Whereas quantification is ultimately desired for human arteries, mouse models of vascular development, health, aging, and disease continue to prove increasingly valuable, particularly given the ability to collect longitudinal data on wild-type, genetically modified, pharmacologically treated, and surgically manipulated mice in large numbers. Given that many normal segments of elastic and muscular arteries are nearly cylindrical, miniaturized biaxial pressure–distension and axial force–extension testing devices have provided considerable data for such quantification.^1–3^ There are many cases, however, wherein the geometry is complex, such as in aneurysms, atherosclerosis, and dissections, which necessitates advanced experimental methods. We thus developed a panoramic digital image correlation (pDIC) method than can be coupled with optical coherence tomographic (OCT) examinations of murine vessels to quantify material properties regionally.^4,5^ This approach has since provided increased insight into diverse conditions, including thoracic aortic aneurysm, abdominal aortic dissection, and aortic tortuosity.^6–8^

Our pDIC approach yields high-fidelity 3D reconstructions of murine vessels imaged in multiple quasi-statically loaded configurations, which allows computation of the full field of local deformations along the vessel surface.^4,9^ To estimate associated best-fit values of constitutive model parameters regionally, we use a novel inverse characterization approach based on the principle of virtual power.^10^ Adding local thickness measurements derived from OCT allows the inverse characterization to be extended from DIC to digital volume correlation, as illustrated in our study of aortic dissections that include significant intramural thrombosis.^6^ Notwithstanding these prior technical advances, there has remained a pressing need for more robust quantification of biomechanical properties, complete with estimates of their associated uncertainties, to enable reliable comparisons of results both across complex lesions and across mouse models. The lack of quantitative uncertainty estimates have made it unfeasible to address questions regarding inferred biomechanical properties on a subject-specific basis, such as whether stiffness differs significantly between multiple material locations on the same vessel, or whether stiffness differs significantly between multiple vessels at a corresponding material location. In previous studies,^5–8^ we have circumvented this issue by performing only group-level analyses of pooled regional results, while also limiting our analysis to material points at which the constitutive model explained a very high proportion of the variance in the observed mechanical data (as quantified by the coefficient of determination). We note, however, that the ability of the model to fit the data well does not necessarily correspond to a high degree of identifiability in the material parameter values, or similarly in derived estimates of the vessel wall’s mechanical properties, such as tangent stiffness or stored energy.^3^ Hence, there was a critical need to extend the modeling framework to provide measures of uncertainty that can be reported alongside the locally estimated material parameters and related quantities of interest.

In the present study, we have integrated a novel uncertainty quantification and propagation pipeline within our inverse modeling approach, relying on empirical and analytic Bayesian techniques. To demonstrate the approach, we present illustrative results for three murine ascending thoracic aorta specimens, quantifying uncertainties in constitutive model parameters as well as circumferential and axial tangent stiffness components (Section 3). We note that consistent comparisons of data across different mouse models promises to provide far greater insight into vascular health and disease (cf. Bellini *et al*. (2017),^11^ Humphrey & Tellides (2019)^12^) than detailed studies that only compare wild-type to a single model of interest, as is common. Toward this end, herein we applied our pipeline to data from three different mouse models: wild-type control, apoliproprotein E-null (*Apoe*^-/-^) infused with angiotensin II (Ang II), and wild-type treated with *β*-aminopropionitrile (BAPN). In addition to facilitating both subject-specific and group-level statistical analyses of the mechanics of the vessel wall, our extended workflow will allow mechanobiological models that utilize estimated tissue-level mechanical properties downstream of the present pipeline to account for their associated uncertainties.

## 2 METHODS

### 2.1 Summary of approach

Herein, we augment previously established procedures for mechanical testing, high-resolution imaging, and inverse modeling with a new integrated workflow for uncertainty quantification and propagation (Figure 1). As before, we first collect standard biaxial mechanical data to examine the gross response of the vessel following established protocols,^3^ which enables estimation of the complete reference configuration (Section 2.2.2). A speckle pattern is then applied to the vessel, which is mounted within a custom device for panoramic imaging. The acquired images are post-processed using pDIC, which yields 3D reconstructions of the vessel in each imaged configuration as well as the complete field of local deformations. Combining the deformation and loading data with local estimates of the wall thickness derived from OCT (Section 2.3) allows inverse modeling to estimate local material model parameters (Section 2.4), from which we can compute mechanobiologically relevant quantities of interest, such as tangent stiffness. To advance this approach, we now quantify uncertainties in the locally estimated model parameters using a Bayesian framework (Section 2.4.1), and then propagate those uncertainties to obtain posterior distributions of our quantities of interest at each point along the vessel wall (Section 2.4.2).

**FIGURE 1.**
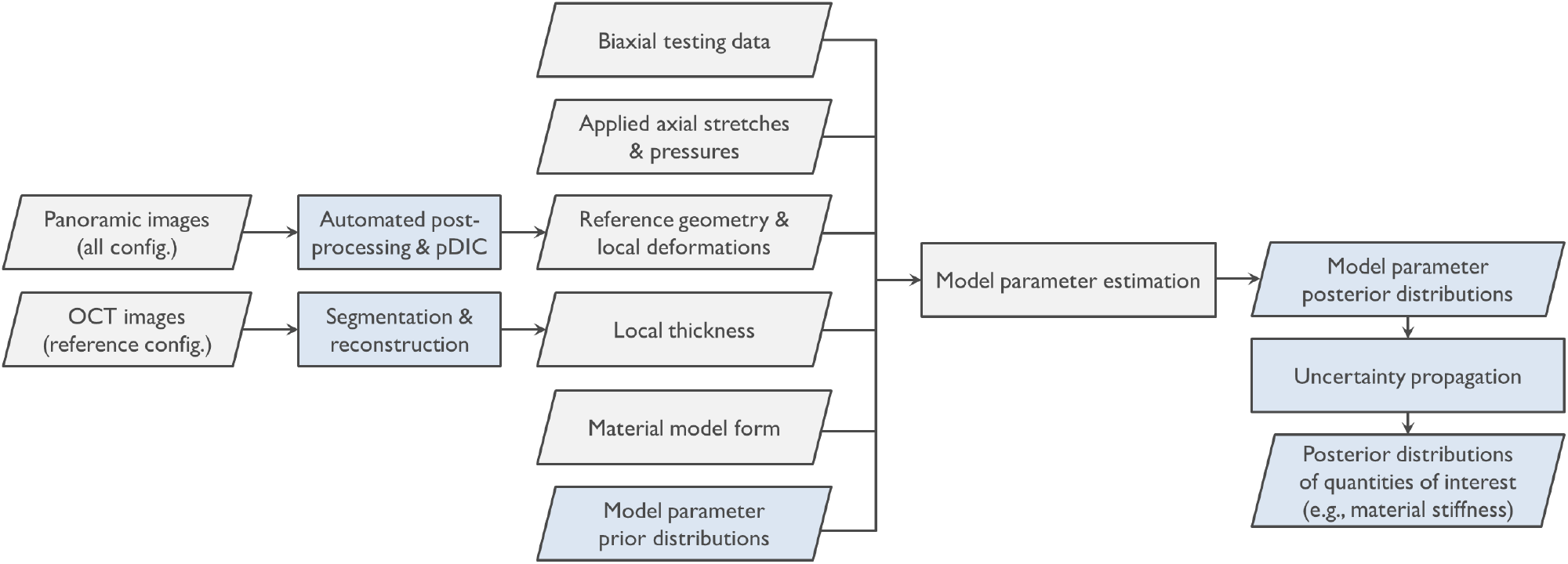
Summary of the complete workflow required to estimate mechanical properties locally. Boxes shown in blue highlight contributions of the present study, which focuses on the quantification of uncertainty in material model parameters as well as the propagation of that uncertainty to broader quantities of interest (e.g., circumferential and axial stiffness) whose estimated values depend on multiple model parameters. config., configuration; OCT, optical coherence tomography; pDIC, panoramic digital image correlation

### 2.2 Data acquisition

#### 2.2.1 Illustrative data set

To demonstrate and validate our novel inverse modeling and uncertainty quantification pipeline, we applied our approach to three ascending thoracic aorta specimens excised from male mice with a C57/BL6 genetic background. To highlight potential differences in the uncertainty of estimated mechanical properties across different specimen types, we herein show results for one wild-type control specimen, one *Apoe*^-/-^ specimen infused with Ang II for 14 days to induce an aneurysm coincident with hypertension,^13^ and one juvenile wild-type specimen treated with BAPN for 14 days to induce a dissection caused by inhibition of collagen cross-linking (Table 1).^14^ All animal protocols were approved by the Yale University Institutional Animal Care and Use Committee and followed methods detailed previously.^3,15^

**TABLE 1.** Illustrative sample characteristics

| Descriptive label | Age | Treatment | Presentation |
| --- | --- | --- | --- |
| Wild-type | 12 wk | — | Normal |
| *Apoe*<sup>-/-</sup> + Ang II | 14 wk | 14 d Ang II infusion | Hypertensive, aneurysmal |
| BAPN | 5 wk | 14 d BAPN oral delivery | Dissected |

#### 2.2.2 Biaxial mechanical testing and panoramic imaging

Noting the crucial importance of biaxial data for the accurate estimation of vessel mechanical properties, we collected gross pressure–diameter and axial force–length data for each vessel following established procedures.^3^ Briefly, each vessel was cannulated, secured with sutures, and mounted within a previously validated computer-controlled testing system that permitted both cyclic distension and axial extension.^16^ Following preconditioning, the vessel diameter was measured at pressures ranging from 10 to 140 mmHg while holding the axial stretch constant at 0.95, 1, and 1.05 times the initially estimated *in vivo* stretch. Similarly, the axial length was measured at axial forces ranging from 0 to 35 mN while holding pressure constant at 10, 60, 100, and 140 mmHg. In preparation for subsequent mechanical data collection within the panoramic imaging system, the estimated stretch value at which axial force would vary the least during pressurization was adopted as an updated estimate of the *in vivo* stretch (*λ*_iv_).^17^

Each vessel was air-brushed with black and white India ink to generate a random speckle pattern over the entire outer surface, then mounted within a panoramic imaging system previously described.^4,9^ Panoramic images were acquired while each vessel was loaded quasi-statically to known values of axial stretch (0.95*λ*_iv_, *λ*_iv_, 1.05*λ*_iv_) and pressure (10–140 mmHg, in steps of 10 mmHg), totaling 42 unique configurations (Figure 2). In each configuration, vessels were imaged from eight views, thus yielding 336 images total per vessel. Due to the high number of images acquired, post-processing of the imaging data for subsequent modeling purposes has historically required about eight hours of interactive input per specimen. In anticipation of the present study and others, we systematically automated this post-processing pipeline, reducing the total time required to less than 30 minutes per specimen with no change in the quality of the results. These automation efforts are described in the Appendix. To arrive at 3D reconstructions of each vessel in every configuration, we performed pDIC between pairs of post-processed images, following previously described methods.^4,9^ Note that the set of 3D reconstructions for each vessel have material point correspondence, enabling the computation of the complete local deformation (tensor) field at every point on the adventitial surface, which is crucial for subsequent inverse modeling and uncertainty quantification (Figure 2; Section 2.4). Each vessel surface was discretized using a 40 × 25 array of quadrilateral elements as in Weiss *et al*. (2020)^8^ prior to inverse modeling.

**FIGURE 2.**
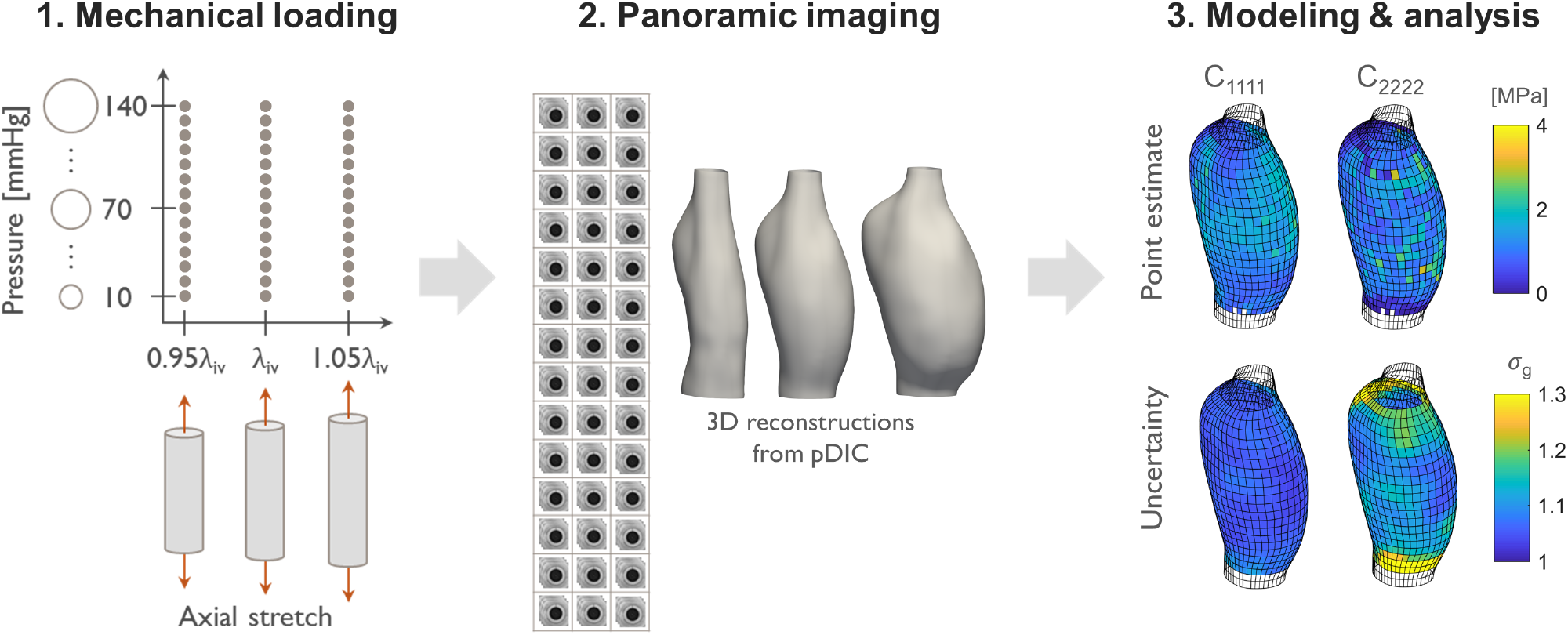
Data acquisition and analysis pipeline. While mounted within the panoramic imaging system, each specimen is loaded to 42 unique quasi-static configurations (3 axial stretches × 14 pressures). In each configuration, specimens are imaged from 8 views to allow complete geometric reconstruction as well as subsequent inverse modeling and analysis. pDIC, panoramic digital image correlation

### 2.3 Local vessel wall thickness estimation

Accurate (local) material model parameter estimation is highly dependent on knowledge of (local) specimen thickness, which is inversely proportional to the estimated stress values used during regression. We recently integrated OCT-derived local thickness measurements within our pDIC-based inverse modeling pipeline, allowing us to no longer rely on simplifying uniform-thickness assumptions.^6^ In the present study, we adopt the approach described in Bersi *et al*. (2019),^6^ but make several practical improvements to the processing of the data as described below. OCT data were collected for each vessel in the reference configuration (70 mmHg pressure, at *in vivo* stretch) at 100 locations along the axial length of the vessel. At each axial location, volumetric images were acquired at four rotationally symmetric positions around the central axis. These four images were registered interactively as in Bersi *et al*. (2019)^6^ to form a complete cross section orthogonal to the central axis (Figure 3).

**FIGURE 3.**
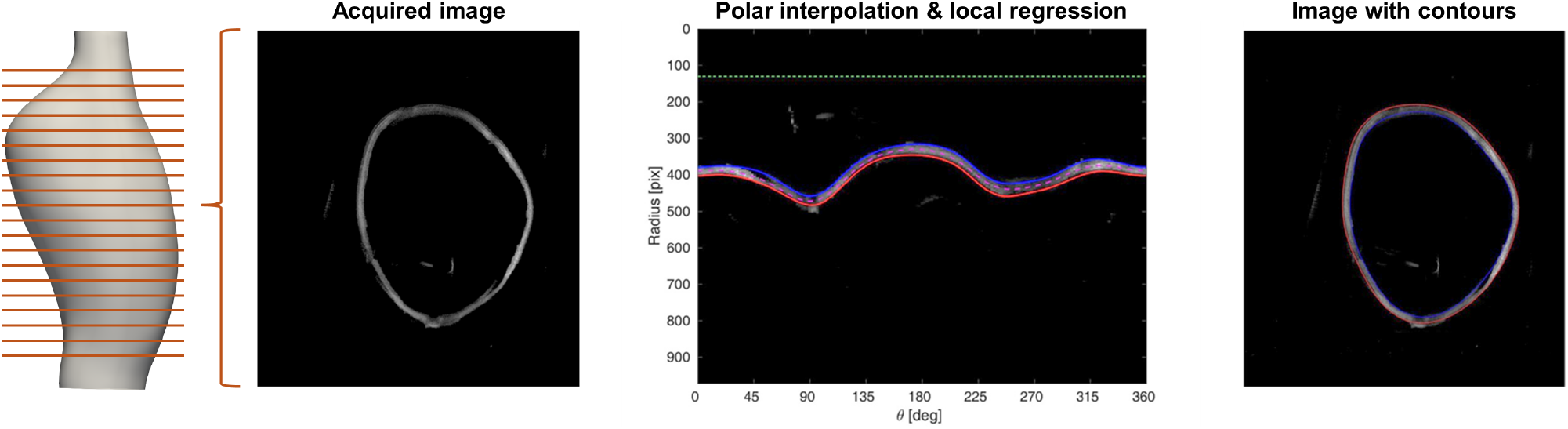
Estimation of local thickness via optical coherence tomography (OCT). Images are acquired at 100 cross-sections (only 20 highlighted above for clarity) along the length of the vessel in its (loaded) reference configuration, and each image is interpolated on a polar coordinate system with the origin at the mean location of all pixels with intensity above a minimum threshold. The medial (magenta), inner (blue), and outer (red) contours of the vessel wall are then estimated using local polynomial regression (LOESS) of high-intensity pixels with radius greater than a minimum value (green)

As an improvement upon previous approaches that required interactive tracing of the inner and outer vessel wall at each cross section, we developed an automated algorithm to extract these contours, thus dramatically reducing the post-processing time as well as eliminating variability resulting from observer subjectivity. First, the image is thresholded using Otsu’s method,^18^ such that high-intensity pixels are categorized as likely tissue regions and low-intensity pixels as likely non-tissue regions. Second, the image data is interpolated on a polar coordinate system, whose origin is at the mean location of all high-intensity pixels (Figure 3). The medial contour of the vessel wall is then obtained via a locally estimated scatterplot smoothing (LOESS) regression of the high-intensity pixel coordinates in polar space.^19,20^ Periodicity was enforced by performing the regression on data which were repeated in the circumferential direction. The local regression model was chosen to be a second-order polynomial with a tricube weight function and span of 90° in the circumferential direction. To remove the outlier effects of spurious bright pixels far from the true vessel wall, only points with radius between 1/3 and 3 times the median radius were included in the regression. The regression was further rendered robust to outliers by performing a second iteration in which the data were weighted using a bisquare function in the radial direction, with a span of 6 times the median absolute radial deviation in the first iteration.^19^

At each circumferential coordinate value *θ_i_*, we computed the minimum and maximum radius residuals, *r*_min,*i*_ and *r*_max,*i*_, from the medial contour. To extract the inner and outer contours, we performed the same LOESS procedure as before, but on the *r*_min,*i*_ and *r*_max *i*_ respectively, and added the result to the medial contour. The local thickness was finally computed as the difference between the inner and outer radius, using the approximation that the surface normal nearly coincides with the radial coordinate direction at all points (this approximation holds except, for example, in the presence of large saccular aneurysms). After the contours of all cross sections are obtained, the 3D outer contour stack is registered to the pDIC-derived reconstruction of the reference configuration using a least-squares–optimal rigid body transformation as described in Bersi *et al*. (2019),^6^ and the local thickness data are mapped to the pDIC-derived geometry using a nearest-neighbor search. Note that the region of the vessel spanned by the OCT data often overlaps only partially with the vessel geometry reconstructed from pDIC. Regions of the vessel wall without reliable thickness data, as well as the ends of the vessel near the sutures, were excluded from subsequent inverse modeling and uncertainty quantification/propagation, and are illustrated as transparent patches in all figures presented herein.

### 2.4 Inverse modeling and uncertainty quantification

#### 2.4.1 Bayesian parameter estimation

In line with several of our previous studies,^3,5,6,21^ we modeled the vessel wall as a four-fiber-family hyperelastic constrained mixture of (1) isotropic elastin, (2) smooth muscle cells (SMCs) and circumferential collagen, (3) axial collagen, and (4) diagonal collagen, with strain energy density of the form

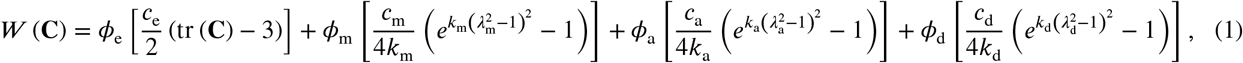

where **C** is the right Cauchy–Green deformation tensor; *ϕ*_(·)_ are mass fractions; *c*_(·)_ and *k*_(·)_ are material parameters; *λ*_(·)_ are direction-specific stretches; and the subscripts (·)_e_, (·)_m_, (·)_a_, and (·)_d_ refer to elastin, circumferential SMCs/collagen, axial collagen, and diagonal collagen, respectively. As in Bellini *et al*. (2014)^21^ and Bersi *et al*. (2016),^5^ stretches *λ*_(·)_ associated with each collagen fiber family were computed from **C** while accounting for the family-specific deposition stretches *G*_(·)_ as well as the in-plane angle *α* of the diagonal family. (We refer the reader to the cited works for background, derivations, and formulæ, which we omit herein for brevity.) At each point on the vessel surface, and for a given set of model parameter values, the vectors of theoretically predicted pairs of pressure and axial force values (**D**_th_ = [**P**_th_, **f**_th_]) can be computed for every configuration using the virtual fields method.^5,10^

It is generally assumed that all of the constituent mass fractions can be accurately prescribed from histology.^5,6^ Therefore, the vector of model parameters remaining to be estimated at each material point is **Ξ** = [*c*_e_, *c*_m_, *c*_a_, *c*_d_, *k*_m_, *k*_a_, *k*_d_, *G*_m_, *G*_a_, *G*_d_, *α*], which are all strictly positive. Adopting a Bayesian framework, we may state our model parameter estimation objective as follows: given the observed mechanical data, and taking into account prior knowledge regarding the model parameter values, find the joint posterior probability distribution of the estimated model parameters. By Bayes’ theorem, the posterior probability density function (PDF) is directly proportional to the product of the likelihood function and the prior PDF.

Without loss of generality, we instead estimate the set of the logarithms of the model parameters, **ξ** = ln(**Ξ**), where ln(·) denotes the natural logarithm; this transformation enables an unconstrained optimization. Moreover, as we describe below, the posterior distribution of **ξ** can be asymptotically approximated as a multivariate normal distribution, which has unbounded support, without violating the theoretical positivity constraints on **Ξ**. The approximate normality of **ξ** also facilitates the analytic approximation of propagated uncertainties (Section 2.4.2).

Following Bersi *et al*. (2016),^5^ we consider the mechanical data to be the paired experimental observations of applied pressure and axial force, **D**_exp_ = [**P**_exp_, **f**_exp_]. To arrive at a point estimate of **ξ**, we formulate the regression problem as a maximization of the posterior probability,

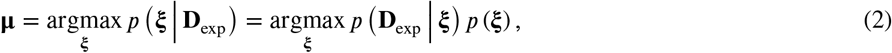

where **μ** is the maximum *a posteriori* estimate of **ξ**, *p*(**ξ** | **D**_exp_) is the posterior PDF of **ξ** given **D**_exp_, *p*(**D**_exp_ | **ξ**) is the likelihood of observing the data given **ξ**, and *p*(**ξ**) is the prior PDF of **ξ**. (Note the shorthand notation *p* for brevity to denote any PDF, though formally these would be symbolically distinguished since they do not have the same form in general. In this shorthand notation, the form of each *p* should thus be inferred from the function argument. In some cases, for clarity, we will distinguish PDFs of different forms using subscripts on *p*·) Equivalently,

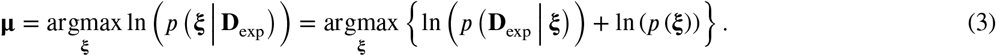

We assume for analytic convenience, as in a least-squares regression, that the residuals **D**_exp,*i*_ – **D**_th,*i*_ are independent and follow a zero-mean multivariate normal distribution with some covariance matrix, which we treat as a nuisance parameter. For simplicity, the prior distribution of **ξ** can also be modeled as a multivariate normal distribution. When there is no prior knowledge, a flat prior may be chosen (*p*(**ξ**) ∝ 1); in this case, the maximum *a posteriori* estimate coincides with the maximum likelihood estimate. Equivalently, the log-prior is chosen to be either quadratic with respect to **ξ** or constant, depending on the availability of prior knowledge. Under these assumptions, it can be shown that at **ξ** = **μ**, the sum of the log-likelihood and logprior (equal to the log-posterior ln(*p*(**ξ** | **D**_exp_)) up to an additive constant) is also asymptotically quadratic with respect to **ξ**; this is known as “Laplace’s approximation.”^22–24^ Since the logarithm of the normal PDF is quadratic, it follows that the posterior distribution of is asymptotically normal, with mean **μ** and covariance matrix 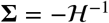, where 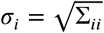 the Hessian matrix of the log-posterior (Figure 4).

**FIGURE 4.**
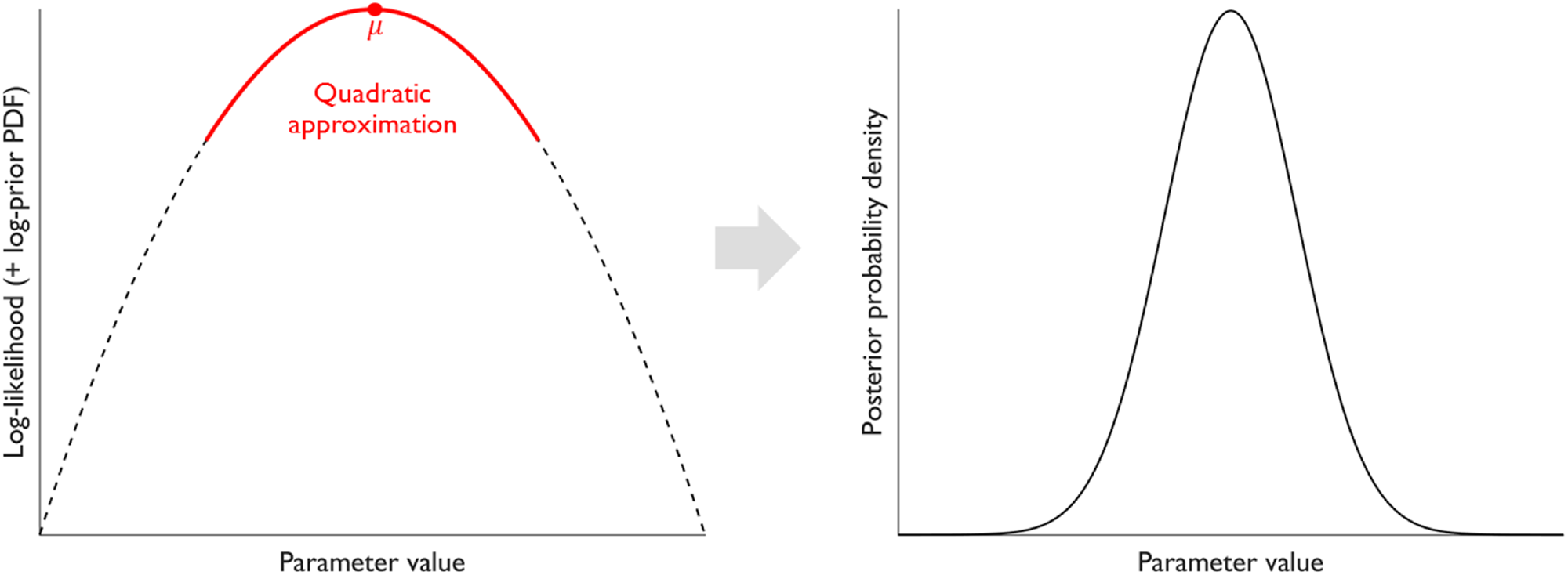
Near a maximum *a posteriori* value *μ*, the logarithm of the posterior probability density function (PDF)—or equivalently, the log-likelihood function alone if a flat prior is used—can be approximated well by a quadratic function (“Laplace’s approximation”), corresponding to a normal approximation of the posterior distribution.^22–24^ For one (multiple) parameter(s), the negative inverse of the curvature (Hessian matrix) corresponds directly to the variance (covariance matrix) of the approximated posterior distribution

Since the posterior distribution of the model log-parameters **ξ** is multivariate normal, the marginal distributions of each *ξ_i_* are also normal, with mean *μ_i_* and standard deviation 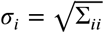. The marginal posterior distributions of the untransformed model parameters Ξ_*i*_ are thus log-normal, with median and geometric mean *μ*_g,*i*_ = *e^μ_i_^*, and geometric standard deviation (GSD) *σ*_g,*i*_ = *e^σ_i_^*, which can be used as a scalar summary of the *relative* uncertainty in a given model parameter. Note that *σ_i_* > 0 ⇒ *σ*_g,*i*,_ > 1, with *σ*_g,*i*_ → 1 denoting complete certainty. From these geometric *distribution* parameters, equi-tailed credible intervals for **Ξ**_*i*_ can be constructed of the form 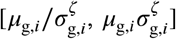, where *ζ* defines the coverage probability (for a 95% credible interval, *ζ* ≈ 1.96). Alternatively, highest posterior density credible intervals can be constructed numerically by searching for the narrowest interval that yields the desired coverage probability.^25^

Using a flat prior, we applied the above inverse modeling framework pointwise along the discretized vessel surface, yielding a set of location-specific posterior distributions for the estimated set of model log-parameters. Due to the cost of computing the gradient of the objective function, we started the optimization by performing 10 iterations using a derivative-free genetic algorithm,^26^ and then used the best current candidate to initialize a local maximizer using the Broyden–Fletcher–Goldfarb– Shanno (BFGS) algorithm,^27,28^ which executed until convergence. The Hessian matrix 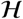 was computed at the optimum **μ** using finite differences.

#### 2.4.2 Uncertainty propagation

Due to the phenomenological nature of the constitutive model, physical interpretability of the specific parameter values is limited, except perhaps in the case of deposition stretches and the angle of the diagonal collagen fiber family. Rather, quantities derived from the evaluation of the constitutive model (conditional on a set of parameter values) are often of primary interest, as they can be related directly to mechanobiological phenomena.^29,30^ These quantities include direction-specific material stiffness, stress components, and stored energy, among others.^8,31^ In addition to uncertainties in the model parameters themselves, it is thus crucial to report uncertainties in these derived quantities of interest, especially for the purposes of model selection, future experimental design, computational (e.g., finite element) simulations, and application studies involving subject-specific and/or tissue-location–specific hypothesis testing. Extending the parameter estimation and uncertainty quantification pipeline presented in Section 2.4.1, we explored two approaches for propagating model parameter uncertainties: Monte Carlo sampling and an analytic approximation based on Taylor series expansion. In Section 3, we present illustrative results for the resulting uncertainties in circumferential (*C*_1111_) and axial (*C*_2222_) tangent stiffness at *in vivo* axial stretch and 100 mmHg of pressure, comparing the two approaches.

While Monte Carlo sampling-based propagation approaches are computationally more expensive, they are asymptotically more accurate, as they rely on weaker assumptions regarding the distribution of the input variables (in this case, the model log-parameters **ξ**).^32,33^ To numerically estimate uncertainties in circumferential and axial stiffness, we randomly sampled 1000 sets of model log-parameters from the posterior distributions of **ξ** at each material point using a Metropolis–Hastings algorithm,^34^ with the proposal density prescribed as a multivariate normal distribution centered at the previous Markov chain sample with covariance **−**. The circumferential and axial components of the tangent stiffness tensor at *in vivo* axial stretch and 100 mmHg of pressure were computed directly as in Bersi *et al*. (2016)^5^ for each random parameter set, and the GSDs *σ*_g_(*C*_1111_) and *σ*_g_(*C*_2222_) were computed from the resulting pointwise samples.

In contrast to sampling-based approaches, analytic approximations to propagated uncertainties have several advantages, including a predefined number of required function evaluations (which can be time-consuming for a computationally complex constitutive model). Further, analytic methods are deterministic, and thus not subject to random variations if computed more than once. Below, we derive a second-order analytic approximation to the propagated uncertainty for the output of an arbitrary nonlinear function of a random vector variable that follows a multivariate normal distribution.

Let 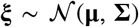 and **δ** = **ξ** – **μ**. Further, let 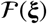 be a nonlinear scalar function of **ξ**, with gradient vector **g** and Hessian matrix **H** at **ξ** = **μ**, such that 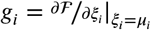 and 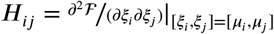. The second-order Taylor series expansion of 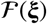 centered at **μ** is then

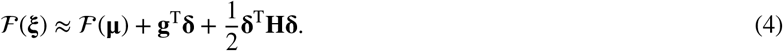

From Equation 4, we can explicitly approximate the expected value of 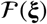 as

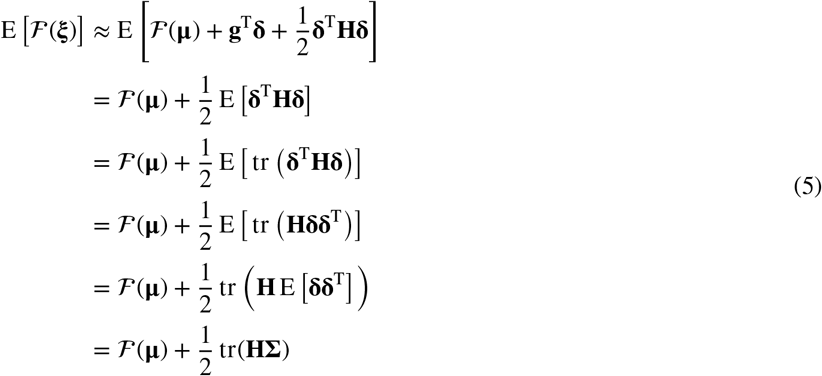

and the variance of 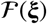 as

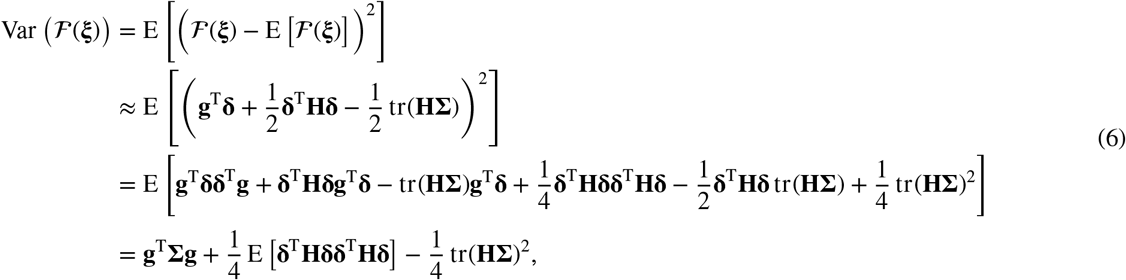

where E[·] denotes the expected value and tr(·) denotes the trace. Note that 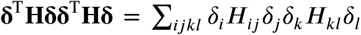. By Isserlis’ theorem,

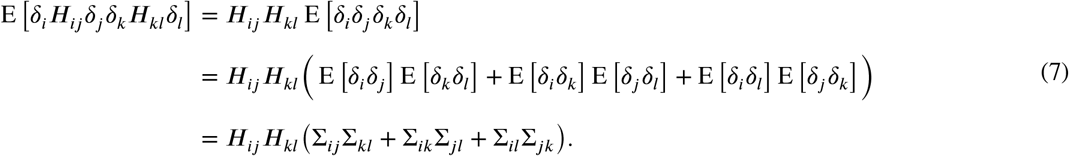

Therefore (exploiting the symmetry of **H** and **∑**),

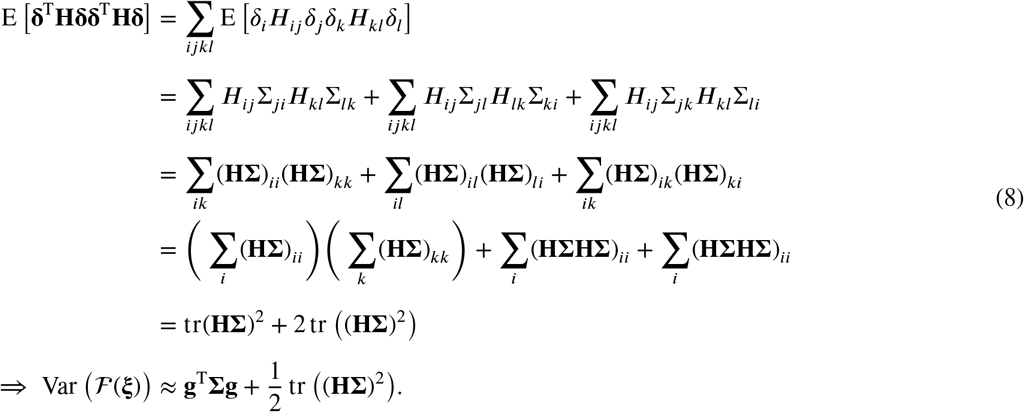

In the case of a vector function 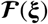, the mean and variance of each component can be independently computed following Equations 6 and 8, while the covariance between components can be similarly derived to yield

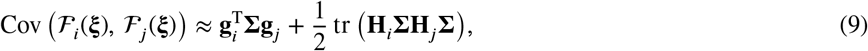

where {**g**_*i*_, **g**_*j*_} are the gradient vectors of 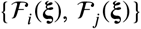 and {**H**_*i*_, **H**_*i*_} are the corresponding Hessian matrices, all evaluated at **ξ** = **μ**.

To apply the above approximation, we first computed the gradient vectors and Hessian matrices numerically using finite differences. The analytic expressions for mean (Equation 6) and variance (Equation 8) were then used directly to compute the pointwise expected value and variance of the logarithm of the circumferential and axial components of the tangent stiffness tensor at *in vivo* axial stretch and 100 mmHg of pressure, from which the GSD of each stiffness component was computed as described in Section 2.4.1.

While the methods described thus far all apply to uncertainty quantification in subject-specific estimates of local mechanical properties, a further extension of our uncertainty propagation framework relates to the computation of posterior distributions of group-level statistics. For example, if pDIC-derived estimates of local stiffness are obtained for several vessels which can be pooled in a meaningful way, what can now be said of the group-wide mean stiffness at a landmark tissue location that is materially correspondent across all specimens? In this case, the propagated uncertainty depends on both the uncertainty of the estimated stiffness in each vessel and the additional uncertainty related to estimating the group-level distribution of estimated stiffness values. To illustrate the propagation of uncertainties to the group level, we derive the posterior distribution of circumferential stiffness at a materially correspondent point *i* across *N* specimens as an example:

Let ln((*C*_1111_)_*i,n*_) denote the circumferential log-stiffness at material point *i* for specimen *n* (note that we again use a logarithmic transformation to map the strictly positive quantity (*C*_1111_)_*i,n*_ to an unbounded space). At the group level, we assume for simplicity that the distribution of log-stiffness is normal, with mean *μ*_group_ and variance 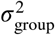. For certain suitable priors on 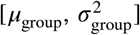, if the true subject-specific ln((*C*_1111_)_*i,n*_) were known exactly, the posterior PDF of the group-level distribution parameters would have a straightforward analytic form. For example, under a normal-inverse-gamma prior, which is conjugate to the normal distribution, the posterior distribution of 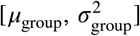 would also be normal-inverse-gamma.^24^ The same is true for the uninformative prior proportional to 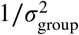, albeit with different distribution parameters. The marginal distribution of *μ*_group_ would be a non-standardized Student’s *t* distribution 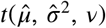 in both of these cases. If the uninformative prior is adopted (reflecting no prior knowledge at the group level), the posterior distribution of *μ*_group_ would be centered at the sample mean 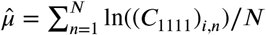 and scaled according to 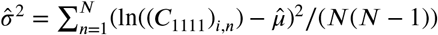, with *v* = *N* – 1 degrees of freedom (for a full derivation, see Zhang *et al*. (2019)^35^).

In practice, of course, the true subject-specific stiffness values are ***unknown*,** which in turn introduces uncertainty in the sample mean and variance that are used to define the posterior distribution of *μ*_group_. Following the Taylor expansion uncertainty propagation approach, we approximate the posterior distribution of ln((*C*_1111_)_*i,n*_) as normal with mean and variance according to Equations 6 and 8. Assuming the specimens are independent, the joint distribution of their log-stiffness values is *N*-variate normal 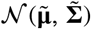, where 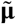 is a vector of the maximum *a posteriori* estimates of ln((*C*_1111_)_*i,n*_) and 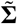 is a diagonal matrix with entries equal to the posterior variances of ln((*C*_1111_)_*i,n*_). The final posterior PDF of the group mean *μ*_group_ is then computed by marginalizing out ln((*C*_1111_)_*i,n*_) using

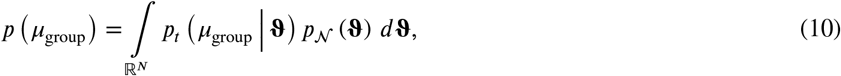

where 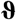 is a dummy variable for integration over the entire domain of ln((*C*_1111_)_*i,n*_), 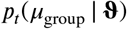 is the PDF of the non-standardized Student’s *t* distribution 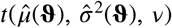, and 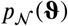 is the PDF of the *N*-variate normal distribution 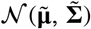. The integral in Equation 10 has no closed-form expression, but is straightforward to compute numerically by quadrature or through Monte Carlo sampling of 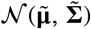, where the result is simply a weighted average of Student’s *t* PDFs. Note that the same approach can be used to compute the bivariate posterior distribution of 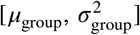, whose PDF 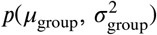 is a weighted average of normal-inverse-gamma PDFs.

While we have outlined the above approach for an individual material point *i*, we note that the analysis could be broadened to an anatomically relevant region of the vessel wall (e.g., anterior, dorsal, etc.). Similarly, for a future specimen *n*^*^, the PDF for the predictive distribution of the subject-specific estimate ln((*C*_1111_)_*i,n*\*_) can be computed by marginalizing out both *μ*_group_ and 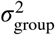 using

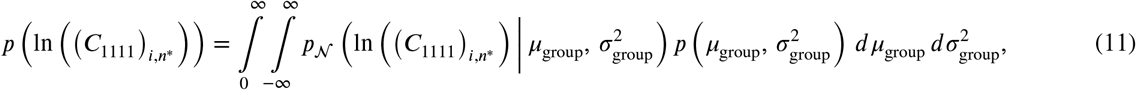

where 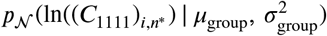 is the PDF of the normal distribution 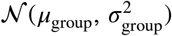.

For both the group-level mean and the new-sample predictive distribution, which are both defined above for the log-stiffness, the PDFs of the corresponding untransformed stiffness values are computed using standard distribution mapping techniques. In the general case, if *Y* = *e^X^*, where *X* is a random “log”-quantity distributed according to a PDF *p*_(*X*)_(*x*), the PDF of *Y* is

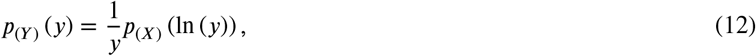

which has strictly positive support.

## 3 ILLUSTRATIVE RESULTS

### 3.1 Uncertainties in material parameters

Application of our novel inverse modeling and uncertainty quantification pipeline to ascending thoracic aorta specimens demonstrated that the resulting uncertainties are heterogeneous across different model parameters as well as throughout different regions of the vessel wall (Figure 5). Notably, uncertainties associated with the material parameters of elastin (*c*_e_) and diagonal collagen (*c*_d_) were substantially higher than those associated with circumferential SMCs/collagen (*c*_m_) and axial collagen (*c*_a_), suggesting that the properties of elastin and diagonal collagen are more difficult to quantify using the examined range of pressure and axial stretch. Nevertheless, GSD values were typically between 1.05 and 1.4, suggesting that in most cases the unobserved/”true” parameter values are close to the maximum *a posteriori* estimate. In the case of a relatively high GSD of 1.4, construction of commonly reported credible intervals indicated that the unobserved material parameter value falls between ~0.5 and 2 times its posterior median with at least 95% probability (Figure 6).

**FIGURE 5.**
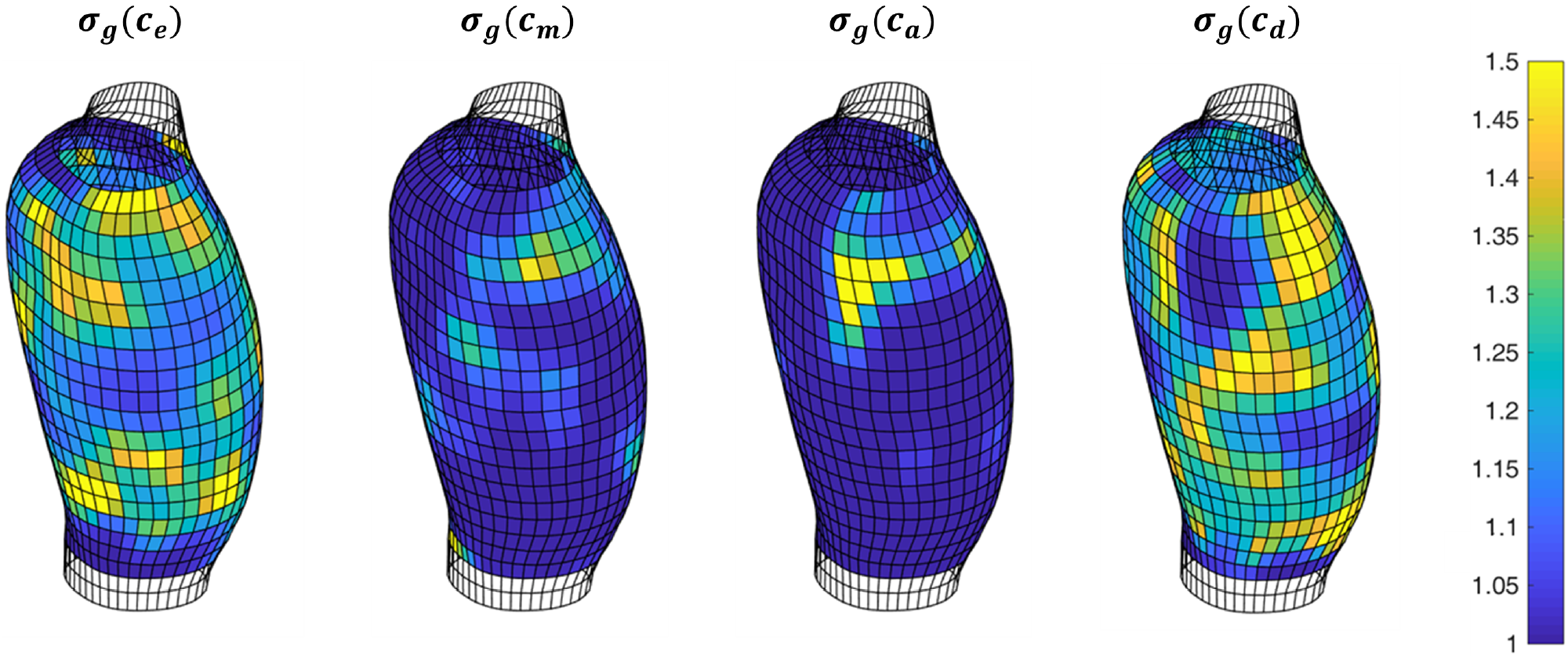
Geometric standard deviation (GSD) maps of material parameters of elastin (*c*_e_), circumferential smooth muscle cells (SMCs)/collagen (*c*_m_), axial collagen (*c*_a_), and diagonal collagen (*c*_d_) for an illustrative wild-type specimen

**FIGURE 6.**
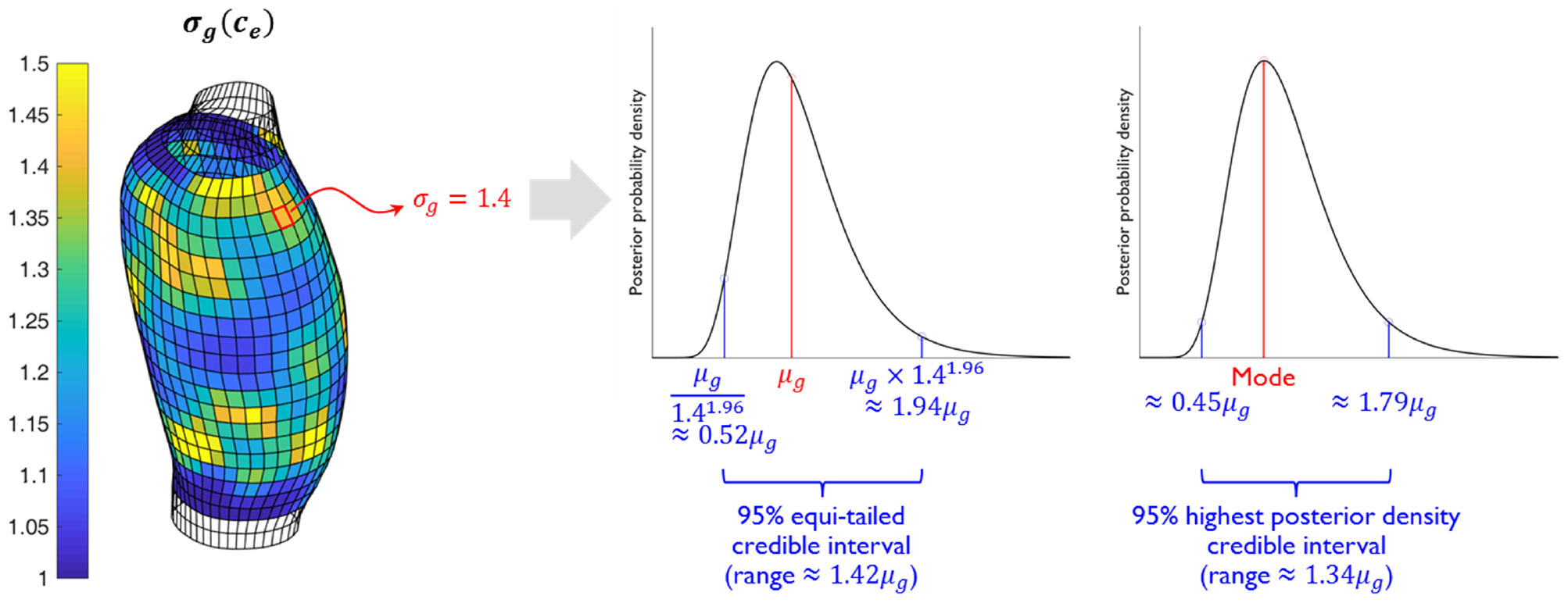
For a strictly positive model parameter (e.g., *c*_e_), the point estimate and geometric standard deviation (GSD) completely define the asymptotically log-normal location-specific posterior distribution, from which credible intervals can be readily derived. For a illustrative location with point estimate (i.e., posterior median) *μ*_g_ and GSD *σ*_g_ = 1.4, 95% equi-tailed and highest posterior density credible intervals are computed as shown

### 3.2 Uncertainties in tangent stiffness components

Propagation of model parameter uncertainties to the circumferential and axial tangent stiffness components at the *in vivo* stretch and 100 mmHg of pressure, using both Monte Carlo sampling and a Taylor series approximation, yielded similar results (Figure 7). At locations on the vessel wall where the two methods differed, the GSD estimated via Monte Carlo sampling was typically higher (by up to ~0.05) than that estimated via Taylor series expansion. This may suggest that the asymptotic assumptions regarding the parameters’ posterior distributions, which are more heavily relied upon in the Taylor approximation, may sometimes lead to an underestimation of the resulting uncertainty–though in general, the two approaches agreed well. Interestingly, the uncertainty in circumferential stiffness (*C*_1111_) was consistently smaller than the uncertainty in axial stiffness (*C*_2222_) along the entire vessel wall (Figure 7); this difference is consistent with the relative paucity of axial stretch states (three, corresponding to three prescribed axial lengths) compared to the circumferential stretch states (14, corresponding to 14 levels of applied pressure) included in the mechanical data that were used for inverse modeling (Figure 2). Moreover, the uncertainties in both stiffness components were often substantially smaller than uncertainties in the individual material parameters. Most notably, the median GSDs of *c*_e_ and *c*_d_ were 1.219 and 1.209, compared to 1.057 and 1.109 for *C*_1111_ and *C*_2222_, respectively. This trend indicates that overall stiffness may be estimated with high credibility even when the model parameters themselves are difficult to estimate; broadly speaking, this is because the quality of the constitutive fit to the mechanical data is influenced jointly by all of the model parameters, whose individual contributions are sometimes weak or partly redundant.

**FIGURE 7.**
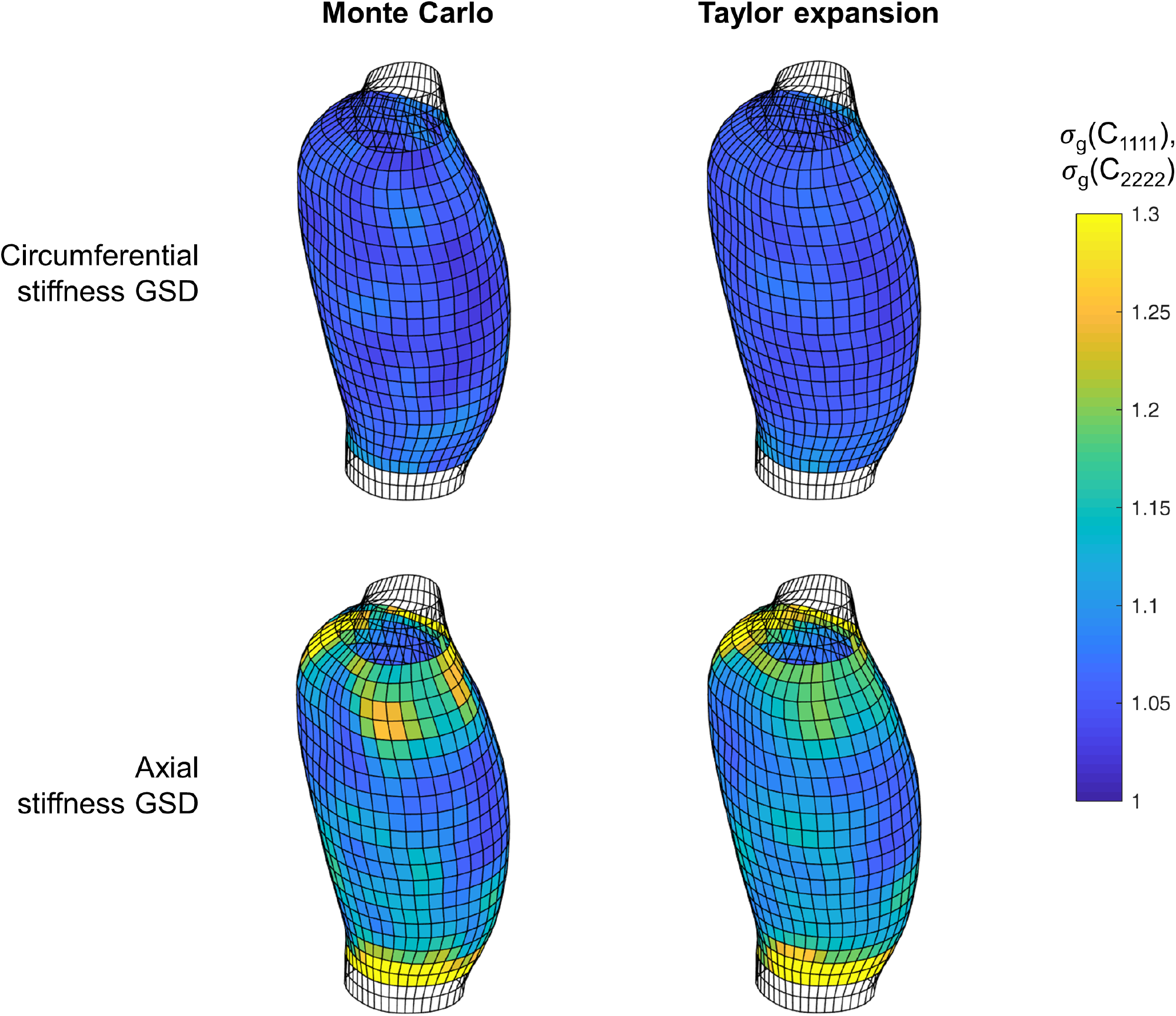
Geometric standard deviation (GSD) maps of circumferential and axial stiffness for an illustrative wild-type specimen, determined by propagating uncertainties in the model parameters through either Monte Carlo sampling or a Taylor series expansion. Under the observed uncertainties, the two methods yield comparable results. Note also that the uncertainties in overall stiffness are often substantially smaller than uncertainties in the individual material parameters (compare to Figure 5, which has different color axis limits)

### 3.3 Comparison between specimen types

We applied our pipeline successfully to both a wild-type control specimen as well as two non-control specimens—one from an *Apoe*-deficient mouse infused with Ang II and one from a wild-type juvenile mouse treated with BAPN (Table 1). The resulting uncertainties associated with circumferential and axial stiffness components varied substantially across specimen type (Figure 8). In general, uncertainties were smallest for the wild-type control specimen, indicating that the model parameters and derived stiffness quantities were more identifiable, despite the constitutive model’s general ability to fit non-control mechanical data well.^6^ The BAPN specimen, which experienced less circumferential stretch during mechanical testing due to its prior dilatation, had notably higher uncertainties in the estimated circumferential stiffness values. Across all specimens, distributions of GSD computed via kernel density estimation^36^ reinforced the finding that uncertainties in axial stiffness were consistently larger than those in circumferential stiffness (Figure 8b).

**FIGURE 8.**
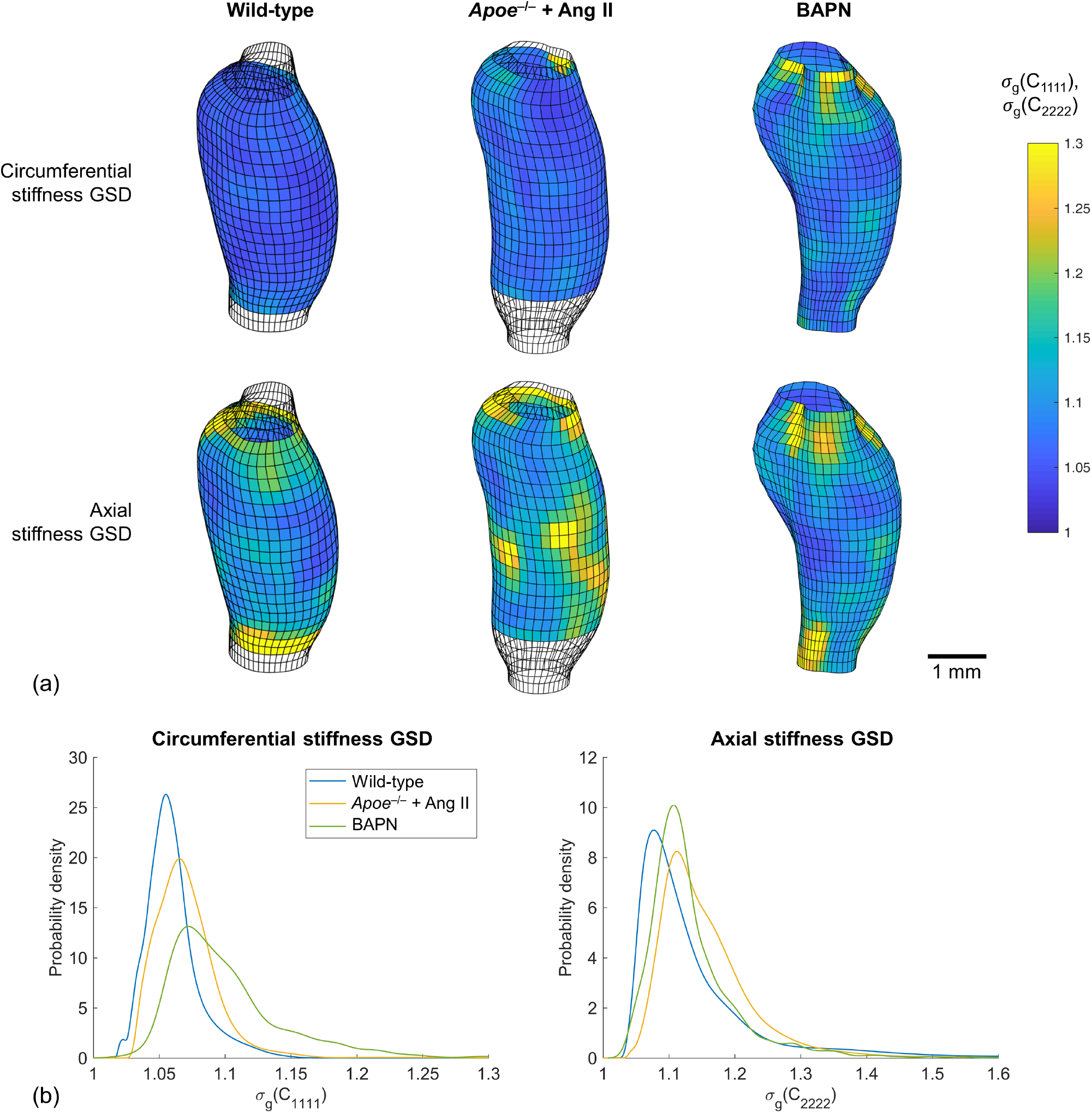
(a) Geometric standard deviation (GSD) maps of circumferential and axial stiffness for the three illustrative specimens, highlighting that the amount of uncertainty can vary substantially across specimen types. All samples are drawn to scale, according to the 1 mm bar shown. (b) Distributions of the GSD of circumferential and axial stiffness over the entire vessel wall for each specimen

### 3.4 Uncertainties at the group level

To demonstrate the propagation of uncertainties to the level of group statistics, we computed the posterior distribution of the group-wide geometric mean of the circumferential stiffness (*μ*_g_((*C*_1111_)_*i*_)) at a materially correspondent point *i* across all three specimens (Figure 9a). Note that we present these results merely to illustrate our uncertainty propagation pipeline, without implying that specimens of these different types would/should be pooled in an application-focused study. Following the assumptions and methods described in Section 2.4.2, and starting from the subject-specific posterior distributions of (*C*_1111_)_*i,n*_ (Figure 9b), the resulting posterior PDF of *μ*_g_((*C*_1111_)_*i*_) is a weighted mixture of non-standardized Student’s *t* PDFs in the log-space mapped to the linear space using Equation 12 (Figure 9c). Similarly, we computed the predictive distribution of new subject-specific estimates of (*C*_1111_)_*i,n*\*_, which is substantially more variable than the group mean (Figure 9d). For the selected point *i*, we found that the shape and effective support of this predictive distribution was similar to the distributions of subject-specific circumferential stiffness estimates over the entire vessel wall (Figure 9e). Broadly speaking, this result indicates that point *i* is not expected to be substantially different from the vessel as a whole.

**FIGURE 9.**
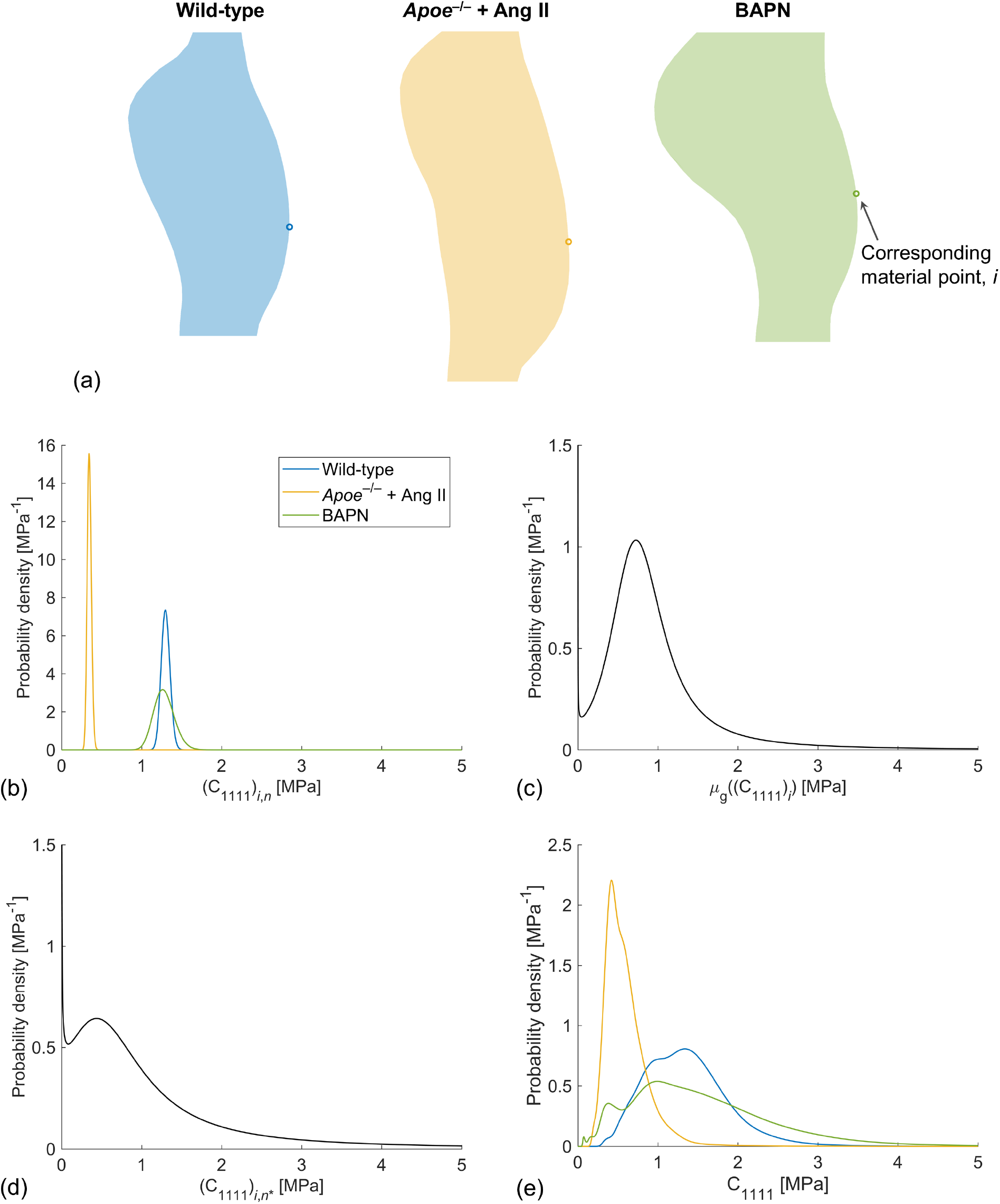
Propagation of uncertainty to group-level statistics. (a) Outlines of the three illustrative specimens, with a subjectively selected corresponding material point *i* highlighted. (b) Posterior distributions of the circumferential stiffness at *i* for each specimen *n*. (c) Resulting posterior distribution of the corresponding group-level geometric mean. (d) Resulting posterior predictive distribution of the circumferential stiffness at *i* for a new (i.e., yet unobserved) specimen *n*^*^. (e) Distributions of circumferential stiffness estimates over the entire vessel wall for each specimen. Note the similar effective support of the distributions in (d) and (e), illustrating that, in this case, the predicted distribution of new stiffness estimates at *i* qualitatively matches the distribution of all previously estimated stiffness values (i.e., broadly speaking, point *i* is not expected to be substantially different from the vessel as a whole)

## 4 DISCUSSION

There is now an extensive literature on nonlinear material properties in health and diverse diseases for arteries from both humans and myriad animal models.^37,38^ In most studies, the associated data and quantification focus on bulk, or homogenized, properties, not site-specific variations. In many cases, however, the nidus of disease is highly localized, with focal differences in material properties nucleating atherosclerotic lesion development, aneurysmal expansion, dissection, and so forth. There is, therefore, a pressing need for more localized descriptions of arterial properties, both within the same lesion and across different examples thereof. Motivated thus, we proposed a pDIC + OCT-based approach for quantifying material properties locally in excised samples subject to *in vivo* relevant mechanical loading. To exploit such information, however, there remained a pressing need both to streamline and automate the data analysis pipeline and to improve the methods of quantification, particularly to assess uncertainty.

In the present study, we have augmented our pDIC + OCT approach to quantify regional material properties with a novel uncertainty quantification and propagation pipeline. We illustrated the utility of this pipeline for ascending thoracic aorta specimens from both a wild-type control mouse as well as two disease models to demonstrate its practical use, and showed illustrative results both for the uncertainties of individual material parameters and the propagation of those uncertainties to tangent stiffness at the subject-specific and group levels. This development represents an important extension of our experimental–computational approach and follows previous milestones in the establishment of our current workflow, including the use of eight panoramic views to ensure high accuracy in pDIC-based reconstructions for strain quantification,^4^ the development of an inverse modeling approach based on the virtual fields method to infer point-wise material parameters,^5^ and the integration of OCT-derived local thickness measurements.^6^ The contribution of the present study makes it possible, for the first time, to assess the identifiability of material parameters (Section 3.1) and the uncertainties in any derived quantities of interest, such as tangent stiffness (Section 3.2), *locally* and on a subject-specific basis. The opportunity to account for subject-specific uncertainties also improves the credibility of group-level statistical analyses (Section 3.4).

While the presented experimental and analytic methods provide a valuable estimate of subject-specific mechanical properties, and especially their spatial heterogeneity as well as associated uncertainties, several practical limitations remain. Most notably, the mechanical testing and image acquisition techniques rely on excised samples, thus precluding longitudinal estimation of mechanical properties for the same subject. While methods to reconstruct the geometry and estimate local deformations of cardiovascular tissues from *in vivo* images are emerging,^7,39,40^ the spatial and temporal resolution provided by current technology is not yet sufficient to eliminate the need for *ex vivo* pDIC- and OCT-based measurements in murine samples. Moreover, estimation of local material parameters directly from *in vivo* images remains difficult, even in human subjects which have a considerably larger vessel geometries.^41–45^

An additional caveat in the application of the presented methods is that the form of the constitutive model is assumed to be true for the purposes of parameter estimation and uncertainty quantification. This assumption carries an important limitation: while the four-fiber-family model used herein is microstructurally motivated, it is nevertheless phenomenological, thus violating the above assumption by definition. (We note that no truly mechanistic models currently exist to describe vascular mechanics, so this limitation is unavoidable.) Nevertheless, using our current model, we have shown that tissue-level quantities of interest such as tangent stiffness can be estimated with fairly high credibility for a variety of specimen types even when the model parameters themselves have higher uncertainties and lack physical interpretation. Caution is warranted in cases of non-pathological (e.g., development) or pathological (e.g., thrombus, calcification) situations wherein the model may have a more limited ability to reproduce the data. Model misspecification introduces an additional layer of uncertainty and potential bias to the analysis, separate from the quantified uncertainties in the parameter values.^46^ It is therefore crucial to confirm that the adopted model has adequately captured the observed mechanical response—for example, by requiring a high coefficient of determination in regions that are included in the analysis,^5^ and by qualitatively verifying independence of the residuals. As an extension of the present work, Bayesian model selection methods can be employed to choose between competing constitutive model forms.^23,47^

## 5 CONCLUSION AND FUTURE WORK

In summary, we integrated a novel uncertainty quantification and propagation pipeline within our inverse modeling approach to estimate local mechanical properties in vascular tissues. The methods rely primarily on well-established empirical and analytic Bayesian techniques, and facilitate both subject-specific and group-level statistical analyses of the mechanical response of the vessel wall. Future directions include the application of the presented pipeline to mechanobiological network-based models of cell signaling as well as predictive models of vascular development, growth, and remodeling.^29–31^ Other potential applications in disease include the comparison of aneurysmal, tortuous, dissected, and/or thrombotic regions of the vessel wall to their normal counterparts.^7,48^ The approach can also be extended by accounting for uncertainties in the acquired data, including histologically prescribed tissue constituent mass fractions, OCT-derived local thickness estimates, and experimental vessel deformations relative to the unloaded configuration.^3,49^

## Nomenclature

Ang II: angiotensin II
ApoE: apolipoprotein E
BAPN: *β*-aminopropionitrile
BFGS: Broyden-Fletcher-Goldfarb-Shanno
GSD: geometric standard deviation
LOESS: locally estimated scatterplot smoothing
OCT: optical coherence tomography
pDIC: panoramic digital image correlation
PDF: probability density function
SMC: smooth muscle cell

## Acknowledgments

This material is based upon work supported by the National Institutes of Health grant no. U01-HL142518.

## Data availability

The data that support the findings of this study are available from the corresponding author upon reasonable request.

## Appendix Automation of panoramic image post-processing

Panoramic images were acquired while each vessel was loaded quasi-statically to known values of axial stretch (0.95*λ*_iv_, *λ*_iv_, 1.05*λ*_iv_) and pressure (10–140 mmHg, in steps of 10 mmHg), totaling 42 unique configurations (Figure 2). In each configuration, vessels were imaged from eight views, thus yielding 336 images total per vessel. Due to the high number of images acquired, post-processing of the imaging data for subsequent modeling purposes has historically required about eight hours of interactive input per specimen. In anticipation of the present study and others, we systematically automated this post-processing pipeline, reducing the total time required to less than 30 minutes per specimen with no change in the quality of the results.

Each panoramic image includes a central nearly circular region displaying the reflection of the vessel wall’s speckle pattern upon a concave conical mirror, surrounded by an array pattern of 245 fiducial markers.^4^ Precise identification of the fiducial marker centroids, as well as registration of the markers between each configuration’s eight acquired images, is crucial to the proper calibration of the imaging system and subsequent mapping of the markers’ image coordinates to real-world positions. Intensity thresholding of the acquired image via Otsu’s method^18^ yields a binarized image in which the fiducial markers as well as most of the tissue region, the corners of the field of view, and spurious dark pixels display as black (Figure A1). At this stage, segmentation of the binarized image into marker versus non-marker regions previously required interactive input through a graphical user interface. We automated the marker identification process by first computing the overall area of each dark region as well as its circularity, which is proportional to the ratio between its area and the square of its perimeter. Since the number of markers greatly exceeded the number of non-marker dark regions, the dark region with the median area and circularity could be classified as a marker with near certainty. Moreover, because all markers had a similar size and shape, all the remaining markers could be expected to have area and circularity close to the median values, relative to the overall range of observed area and circularity values. Each dark region was therefore categorized as a marker if its [circularity, area] vector was one of the 245 nearest neighbors to the median [circularity, area], as measured by the Mahalanobis distance,^50^ which is computed using the standardized (i.e., normalized by standard deviation) circularity and area values. To establish correspondence between images from different views, the markers were then automatically sorted based on their azimuthal angle and distance from the mean of the marker centroid locations.

**FIGURE A1.**
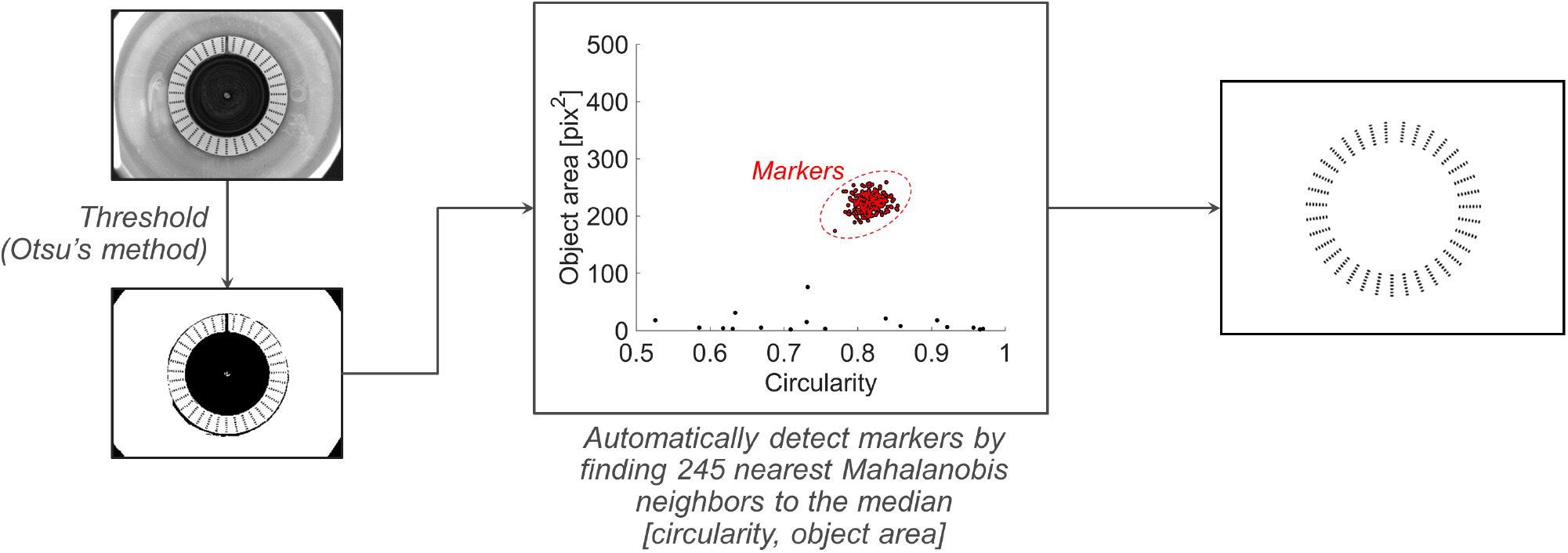
Automatic identification of fiducial markers in panoramic images. An original acquired image is binarized using an intensity threshold determined via Otsu’s method,^18^ and the area and circularity of each dark region is computed. Then, each dark region is categorized as a marker if its [circularity, area] vector is one of the 245 nearest neighbors to the median [circularity, area], as measured by the Mahalanobis distance (identified markers are highlighted in red)

The correlation-based local image registration methods which are the foundation of pDIC depend on a high degree of similarity in the horizontal and vertical directions between the two images of interest; the acquired panoramic images, however, which correspond to eight radially symmetric camera positions, exhibit circumferential and radial similarity instead. Therefore, as in previous works by Genovese *et al*.^4,9^ and Bersi *et al*.,^5,6^ pDIC was performed not using the acquired images directly, but instead on “rectified”/“unwrapped” images; that is, an interpolation of the original image data on a polar coordinate system. Because the camera is not exactly coaxial with the vessel and conical mirror being imaged, the center point of the acquired image does not correspond in general with the vertex of the conical mirror about which the image data should be unwrapped. Unwrapping the image data about a point far from the conical vertex yields characteristic undulation patterns in the interpolated image data (Figure A2), which can cause pDIC to perform poorly. Previously, the search for an acceptable unwrapping center point relied on several iterations of interactive guessing, until an unwrapped image with minimal distortions was obtained; this process was required to be performed for every image acquired (336 images total per vessel, if the examined configurations correspond to the protocols described in Section 2.2.2).

For the present study, we automated this procedure using the following approach: First, the image data is unwrapped using the center of the image as an initial guess of the true center of the imaging setup. Under an optimal unwrapping, the inner boundary of the fiducial marker strip, which is nearly circular in the original image, would map to a nearly straight and horizontal line in polar space. The location of the marker strip boundary can be detected column-wise in the unwrapped image using an intensity threshold, since the marker strip is much brighter than the tissue region (Figure A2). Due to the inaccuracy of the initial guess, the resulting unwrapped marker strip exhibits the undulation pattern described above, where the vertical coordinate of the strip boundary corresponds to the radius coordinate of a circle not centered at the origin. Specifically,

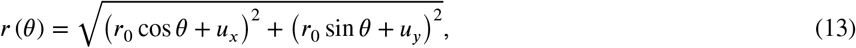

where *r* is the radius coordinate in the (sub-optimally) unwrapped image, *θ* is the angular coordinate, *r*_0_ is the true constant inner radius of the marker strip, and [*u_x_, u_y_*] is the discrepancy between the true center point and the initial guess. Using this relation, the optimal center point can be identified in one iteration by performing a nonlinear least-squares regression of the marker strip radius curve with respect to *r*_0_, *u_x_* and *u_y_* (Figure A2). The original image is then unwrapped with respect to the optimal center point, so that it may be precisely registered to corresponding images from different views and tissue configurations using pDIC.

Using methods described in detail by Genovese *et al*.,^4,9^ we used the pDIC-based image registration results to solve for the real-world 3D position of each point in the acquired images, which yielded a point cloud that represented the vessel wall geometry with high fidelity (Figure A3a). Computation of the local deformation gradient field requires a surface representation of the vessel wall, however, which must be smooth to allow for differentiation of the deformed material point coordinates with respect to their coordinates in the reference configuration. Previously, the point cloud representation was converted to a surface through the use of regression using non-uniform rational basis splines, with the final result depending on subjectively selected shape parameters to compromise between surface smoothness and deviation from the point cloud data. To remove the need for user input, we developed the following automated procedure, which yields a fully parametrized smooth surface reconstruction of the vessel wall in the reference configuration (*in vivo* axial stretch under 70 mmHg pressure): First, the vessel centerline is approximated by applying a Gaussian filter to the *x* and *y* point cloud data in the *z* direction (i.e., along the central axis of the vessel), with the filter span equal to 5% of the total vessel length. Each point is then parametrized with respect to a polar cylindrical coordinate system centered locally at the corresponding centerline location (Figure A3a). To arrive at a smooth surface representation, the local radius coordinate field is modeled as a Gaussian process,^51^ with *θ* and *z* as predictor variables (Figure A3b). Regression is performed on data that are repeated in the *θ* direction, to enforce periodicity. The reconstruction and parametrization is completed by mapping the Gaussian process representation of the local radius coordinate field back to Cartesian space (Figure A3c). To transform the surface reconstruction of the reference configuration to all other configurations, the pointwise *x*, *y*, and *z* displacement fields (computed using the point cloud data) were similarly modeled as Gaussian processes, and the resultant smooth displacement fields were applied to the reference geometry.

**FIGURE A2.**
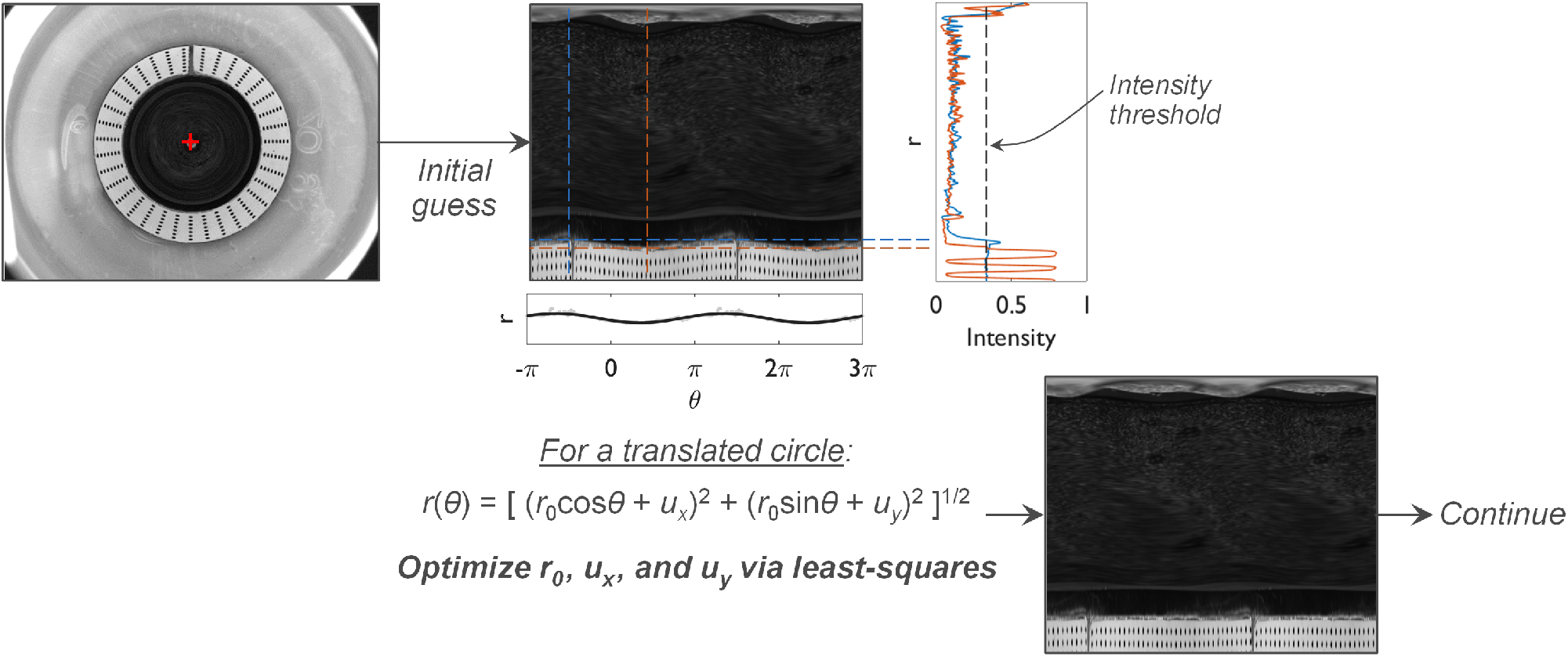
Automatic polar parametrization of panoramic images. For each panoramic image, the image data is interpolated on a polar coordinate system (“unwrapped”) using the center of the image as an initial guess of the true center of the imaging setup. Inaccuracies in the center point about which the image is unwrapped will cause obvious undulations in the array of fiducial markers used to map the image data to real-world coordinates. At each angular coordinate value *θ_i_* in the polar image, the radius *r_i_* of the boundary of the fiducial marker strip can be identified using an intensity threshold. Since the strip boundary *r*(*θ*) will closely match the radius of a translated circle (i.e., one whose center is not the origin), an optimal correction vector [*u_x_, u_y_*] to the initially guessed center point can be determined via least-squares regression. The original image is then unwrapped with respect to the optimal center point, so that it may be precisely registered to corresponding images from different views and tissue configurations

**FIGURE A3.**
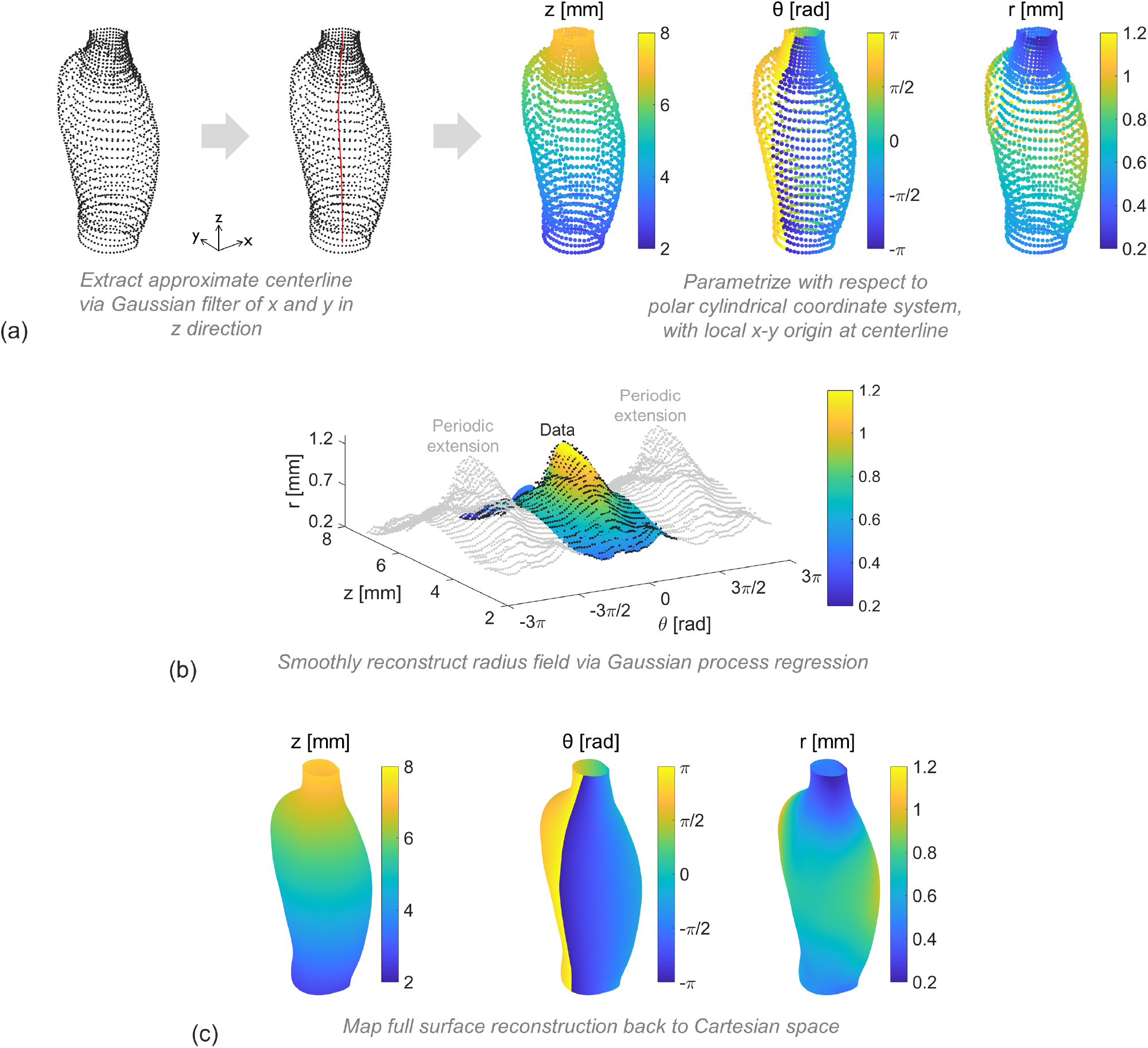
Smooth surface reconstruction of the entire vessel wall. (a) Starting from a panoramic digital image correlation (pDIC)-derived point cloud of the vessel wall, the vessel centerline is approximated by applying a Gaussian filter in the *z* direction to the *x* and *y* data. Each point is then parametrized with respect to a polar cylindrical coordinate system centered locally at the corresponding centerline location. (b) To arrive at a smooth surface representation, the local radius coordinate field is modeled as a Gaussian process, with *θ* and *z* as predictor variables. Regression is performed on data that are repeated in the *θ* direction, to enforce periodicity. (c) The reconstruction and parametrization is completed by mapping the Gaussian process representation back to Cartesian space

